# Cryo-EM structures reveal how ATP and DNA binding in MutS coordinate the sequential steps of DNA mismatch repair

**DOI:** 10.1101/2021.06.03.446775

**Authors:** Alessandro Borsellini, Vladislav Kunetsky, Peter Friedhoff, Meindert H. Lamers

**Affiliations:** Department of Cell and Chemical Biology, Leiden University Medical Center, Leiden, The Netherlands; Institute for Biochemistry, Justus-Liebig University, Giessen, Germany

## Abstract

DNA mismatch repair detects and removes mismatches from DNA reducing the error rate of DNA replication a 100-1000 fold. The MutS protein is one of the key players that scans for mismatches and coordinates the repair cascade. During this, MutS undergoes multiple conformational changes that initiate the subsequent steps, in response to ATP binding, hydrolysis, and release. How ATP induces the different conformations in MutS is not well understood. Here we present four cryo-EM structures of *Escherichia coli* MutS at sequential stages of the ATP hydrolysis cycle. These structures reveal how ATP binding and hydrolysis induces a closing and opening of the MutS dimer, respectively. Additional biophysical analysis furthermore explains how DNA binding modulates the ATPase cycle by preventing hydrolysis during scanning and mismatch binding, while preventing ADP release in the sliding clamp state. Nucleotide release is achieved when MutS encounters single stranded DNA that is produced during the removal of the daughter strand. This way, the combination of the ATP binding and hydrolysis and its modulation by DNA enable MutS to adopt different conformations needed to coordinate the sequential steps of the mismatch repair cascade.

## Introduction

DNA mismatch repair is an evolutionary conserved mechanism that removes mispaired bases from the DNA after DNA replication. Doing so, it reduces the mutation frequency a 100 to a 1000-fold, preventing cancer and drug resistance. It also plays roles in regulation of recombination, triplet repeat expansion and DNA damage signalling ^1,2^. The repair process consists of a cascade of proteins that act sequentially. The MutS protein scans the DNA for mismatches, and upon recognition recruits a second protein MutL. In *Escherichia coli* (*E. coli*), MutL subsequently activates the endonuclease MutH that creates a nick in the newly synthesized strand on hemi-methylated GATC sites ^3–5^. In other species, including eukaryotes, the endonuclease activity resides in the MutL homologs themselves ^6,7^ and is directed to the newly synthesized strand by the DNA sliding clamp (β or PCNA) ^8,9^. In bacteria, the single stranded nick then forms the entry point for the UvrD helicase, or RecD2 in some Bacillus species ^10,11^, that with the aid of a 5’-3’ or 3’-5’ exonuclease will excise the newly synthesized strand ^12,13^. The removed stretch of DNA will then be resynthesized by a DNA polymerase and the remaining nick sealed by a DNA ligase ^14^. In the eukaryotic system the strand removal process is less well understood but involves the replicative DNA polymerase δ and exonuclease EXO1 ^15^.

Throughout this process, MutS is the master coordinator that initiates the subsequent steps of the repair cascade. First, it scans the genome in search for a mismatch. Once it is bound to a mismatch it undergoes a conformational change into a sliding clamp that recruits the second protein MutL ^16,17^. In the clamp state, MutS remains on the DNA for prolonged times and only releases the DNA when it encounters a DNA ends ^18,19^ or a ssDNA gap ^20,21^. This way, MutS molecules will remain present on the DNA until the mismatch has been removed and the repair process is terminated. Recently we have determined multiple cryo-EM structures of MutS on DNA at different stages of the repair process, revealing the multiple conformations that enable it to perform its different tasks ^22^. ATP binding and hydrolysis are essential for MutS to switch between the different states ^23^, but the molecular basis of these transitions and how they are regulated remains unresolved. To gain insight into the ATP-induced conformational changes in MutS we determined four structures of apo MutS at sequential stages of the ATP hydrolysis cycle: ADP-bound, ATP-bound, bound to a non-hydrolysable ATP analogue AMPPNP and bound to the transition state analogue ADP-vanadate ^24^ (ADP-VO_4_^3-^ / ADP-Vi). These structures show how an open MutS dimer in the ADP-bound form closes upon binding to a single ATP but does not reach a hydrolysis competent active site in this structure. Only upon binding of two triphosphate nucleotides the complete active site is formed that can hydrolyse ATP.

Additional biochemical analysis combined with the recently determined structures of DNA-bound MutS, explains how ATP and DNA binding work together to initiate the subsequent steps of the repair cascade. During both the scanning of the DNA as well as mismatch binding, the DNA keeps the ATPase domains in an open, non-catalytic state where the nucleotide binding sites are free to exchange ADP for ATP. Two ATP molecules are needed to transform MutS into the sliding clamp conformation that recruits MutL. In this clamp state, the ATPase active sites are completed and can hydrolyse ATP. However, the dsDNA itself keeps the nucleotide binding sites closed and prevents nucleotide release until MutS reaches a stretch of ssDNA where the interactions with the dsDNA are lost. This enables the dimer to open up and reset the ATPase site for a new round of ATP and DNA binding.

## Results

### Structures apo MutS at sequential stages of the ATP hydrolysis cycle

In order to gain insight into the mechanism of ATP hydrolysis in MutS, we determined four cryo-EM structures of the MutS dimer in absence of DNA at sequential stages of the hydrolysis cycle: 1) bound to two ADP molecules, 2) bound to one ADP and one ATP, 3) bound to two AMPPNP molecules, a non-hydrolysable ATP analogue, and 4) bound to two ADP-vanadate (ADP-Vi) molecules, a transition state analogue (Fig. 1, Table 1, and Extended Data Figs. 1–4). The four structures were determined to a resolution of 4.8, 3.3, 3.4, and 3.7 Å, respectively. All structures were obtained by incubating 4 µM MutS with 3 mM of the nucleotide for 5 min at 4 °C in buffer containing Mg^2+^ before plunge freezing in liquid ethane. In the resulting cryo-EM data, we find that three of the four conditions show a single conformational state: an open form for ADP_2_ and a closed form for both AMPPNP_2_ and ADP-Vi_2_. In contrast, in the sample incubated with ATP we observe both the open ADP_2_-bound form and the closed ATP:ADP-bound form indicating that under these conditions MutS can hydrolyse ATP.

**Fig. 1.**
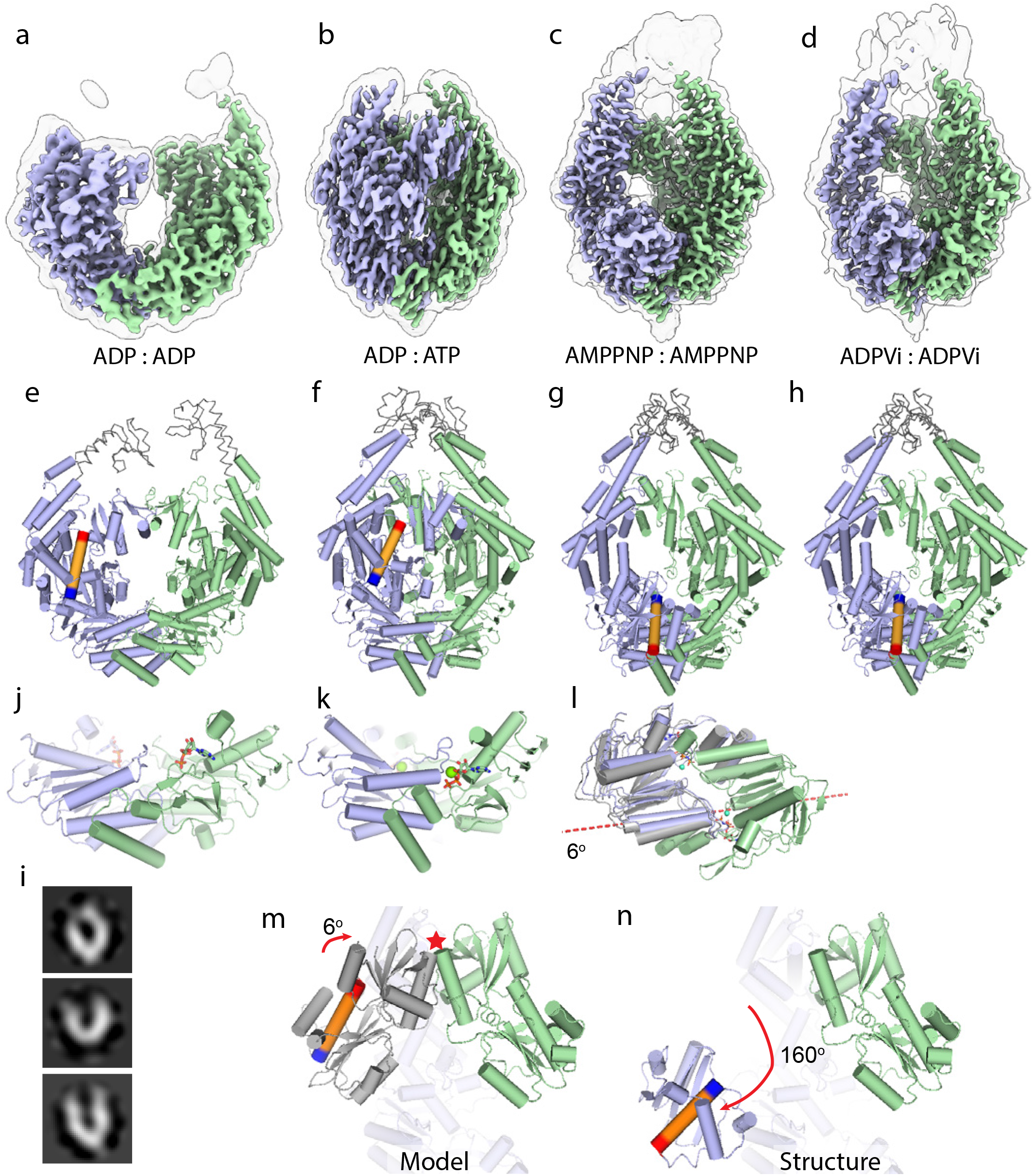
Structures of MutS at sequential steps of the ATP hydrolysis cycle. Cryo-EM maps of **a**, ADP-bound MutS, **b**, ATP-bound MutS, **c**, AMPPNP-bound MutS and **d**, ADP-Vi-bound MutS. Monomer A is shown in green, monomer B in blue. Transparent outlines show the same cryo-EM maps at lower contour level. **e-h**, Models of ADP, ATP, AMPPNP, and ADP-Vi bound MutS. Grey ribbon part of the models indicate approximate position of the missing clamp and lever domains that are poorly defined in the cryo-EM map. A central helix in the connector domain (residues 231 to 248) is colored in orange, with N- and C-termini colored in blue and red, respectivily. **i**, Negative stain 2D classes of ADP-bound MutS showing the flexibility of the MutS dimer. **j-k**, Transition of the ATPase domains from ADP_2_ to ATP:ADP-bound MutS that is a result of a 35° rotation of the two monomers. Yellow arrows indicate helix dipole that is attracted by the negative charge of the triphosphate in the opposite monomer. **k**, Transition of the ATPase domains from the ATP:ADP to AMPPNP_2_-bound MutS. Grey model represent the original position of monomer B before the 6° rotation, dashed red line represents the rotation axis of monomer B. **m**, Model of the AMP-PNP_2_-bound MutS without the movement of the mismatch and connector domain. The red star indicated the clash the between mismatch binding domain of monomer A (coloured green) and monomer B (coloured grey). Core domain and ATPasee domain omitted for clarity. **n**, The rotation of mismatch and connector domain circumvents the clash with monomer A.

**Table 1.** Cryo-EM data collection, refinement and validation statistics.

| **Table 1 \| Cryo-EM data collection, refinement and validation statistics** |  |  |  |  |
| --- | --- | --- | --- | --- |
|  | ADP | ATP:ADP | AMPPNP | ADP-Vi |
| **Data collection and processing** |  |  |  |  |
| Magnification | x130,000 | x105,000 | x130,000 | x130,000 |
| Voltage (kV) | 300 | 300 | 300 | 300 |
| Electron exposure e<sup>-</sup>/Å<sup>2</sup> | 57 | 54 | 54 | 34 |
| Defocus range (μm) | -1 to -2.5 | -0.5 to -2 | -1 to -2.5 | -0.5 to -2 |
| Pixel size (Å) | 1.06 | 0.866 | 1.06 | 0.66 |
| Symmetry imposed | C1 | C1 | C1 | C1 |
| Initial particle images (no) | 110,730 | 527,979 | 294,231 | 496,416 |
| Final particle images (no) | 41,730 | 151,672 | 147,899 | 81,316 |
| Map resolution (Å) | 4.8 | 3.3 | 3.4 | 3.8 |
| FSC threshold | 1.43 | 1.43 | 1.43 | 1.43 |
| Map resolution range (Å) | 4.5 to >10.7 | 3.3 to >4.8 | 3.4 to >4.3 | 3.7 to >6.9 |
| **Refinement** |  |  |  |  |
| Initial model used (PDB code) | 1E3M | AMPPNP | 1E3M | AMPPNP |
| Model resolution (Å) | 7.8 | 3.6 | 3.4 | 4.0 |
| FSC threshold | 0.5 | 0.5 | 0.5 | 0.5 |
| Map sharpening B factor (Å<sup>2</sup>) | -50 | -80 | -80 | -70 |
| **Model comparison** |  |  |  |  |
| Nonhydrogen atoms | 10338 | 10955 | 10336 | 10800 |
| Protein residues | 1320 | 1393 | 1318 | 1373 |
| **B factors (Å<sup>2</sup>)** |  |  |  |  |
| Protein | 35-159 | 35-337 | 34-307 | 21-306 |
| **r.m.s deviations** |  |  |  |  |
| Bond lengths (Å<sup>2</sup>) | 0.013 | 0.012 | 0.014 | 0.013 |
| Bond angles (°) | 1.552 | 1.492 | 1.783 | 1.627 |
| **Validation** |  |  |  |  |
| MolProbity score | 1.20 | 1.31 | 1.16 | 1.26 |
| Clashscore | 4.19 | 5.73 | 3.71 | 4.94 |
| Poor rotamers (%) | 0.09 | 0.96 | 0.00 | 0.09 |
| **Ramachandran plot** |  |  |  |  |
| Favored (%) | 98.46 | 98.39 | 98.46 | 97.64 |
| Allowed (%) | 1.46 | 1.61 | 1.46 | 2.36 |
| Disallowed (%) | 0.08 | 0.00 | 0.08 | 0.00 |

In all four structures, the clamp domains are poorly resolved, indicating that in the absence of DNA these domains are flexible. The flexibility is most pronounced in the structure of ADP-bound MutS, where the lever and clamp domains adopt different conformations that can be discerned by negative stain electron microscopy (Fig 1i). The four structures show an increasing compaction going from the open form in the ADP_2_-bound state, to ATP:ADP-bound state, to the most compact form in the AMPPNP_2_ and ADP-Vi_2_-bound state that both adopt an identical conformation. The transition from the ADP_2_-bound to ATP:ADP-bound is characterized by a 35° rotation of the two monomers towards each other around a pivot point located at the interface of the two ATPase domains (Fig. 1j-k, Supplementary Video 1). It is noteworthy that the addition of a single PO_4_^3-^ ion (from ADP to ATP) results in a dramatic increase in resolution, from 4.8 to 3.3 Å, due to the more stable conformation MutS adopts in the ATP-bound form. The ATP:ADP to AMPPNP_2_/ADP-Vi_2_ transition is the result of an additional ∼6° twist of the ATPase domains along the axis of the helix at the bottom of the ATPase domain (Fig. 1l), resulting in a further closing of the nucleotide binding site (Supplementary Video 1-2). This second rotation would result in a clash of the mismatch-binding domains of the two monomers (Fig. 1m), which may be the reason that the connector- and mismatch domains in one of the two monomers rotate towards the ATPase domains (Fig. 1n, Supplementary Video 3). This rotation is identical to that observed in the ATP-induced DNA-bound, clamp state MutS ^22,25^. However, in clamp state MutS both monomers have rotated the connector and mismatch domains. Of the four structures, only the ADP-bound state is compatible with DNA binding, as the closing of the lever domains in the ATP and AMPPNP/ADP-Vi bound structures blocks the entry channel for the DNA. This suggests that both monomers need to have hydrolysed ATP before MutS can bind DNA.

### Closing of the MutS dimer completes the ATPase active site for hydrolysis

Like all ABC ATPases, the MutS ATPase domains are composite active sites where residues from both monomers are required for ATP hydrolysis ^26^. The four structures presented in this work illustrate how the two ATPase domains of both monomers come together to complete the MutS active site during ATP hydrolysis (Fig. 2a). The completion of the ATPase active site is driven by the negative charge of the γ-phosphate in ATP that forms an attractive force to the positive charge of the helix dipole ^27^ that is deposited via a serine (Ser668 in MutS) in the opposing monomer ^28^ (Fig. 1j-k, 2f). In the open, ADP_2_-bound form, the nucleotide is exposed and free to exchange with ATP molecules from solution (Fig. 2b). In contrast, in the closed ATP:ADP-bound and AMPPNP_2_/ADP-Vi_2_-bound structures the nucleotides in both monomers are encapsulated and likely slowed down in leaving the active site (Fig. 2c).

**Fig. 2.**
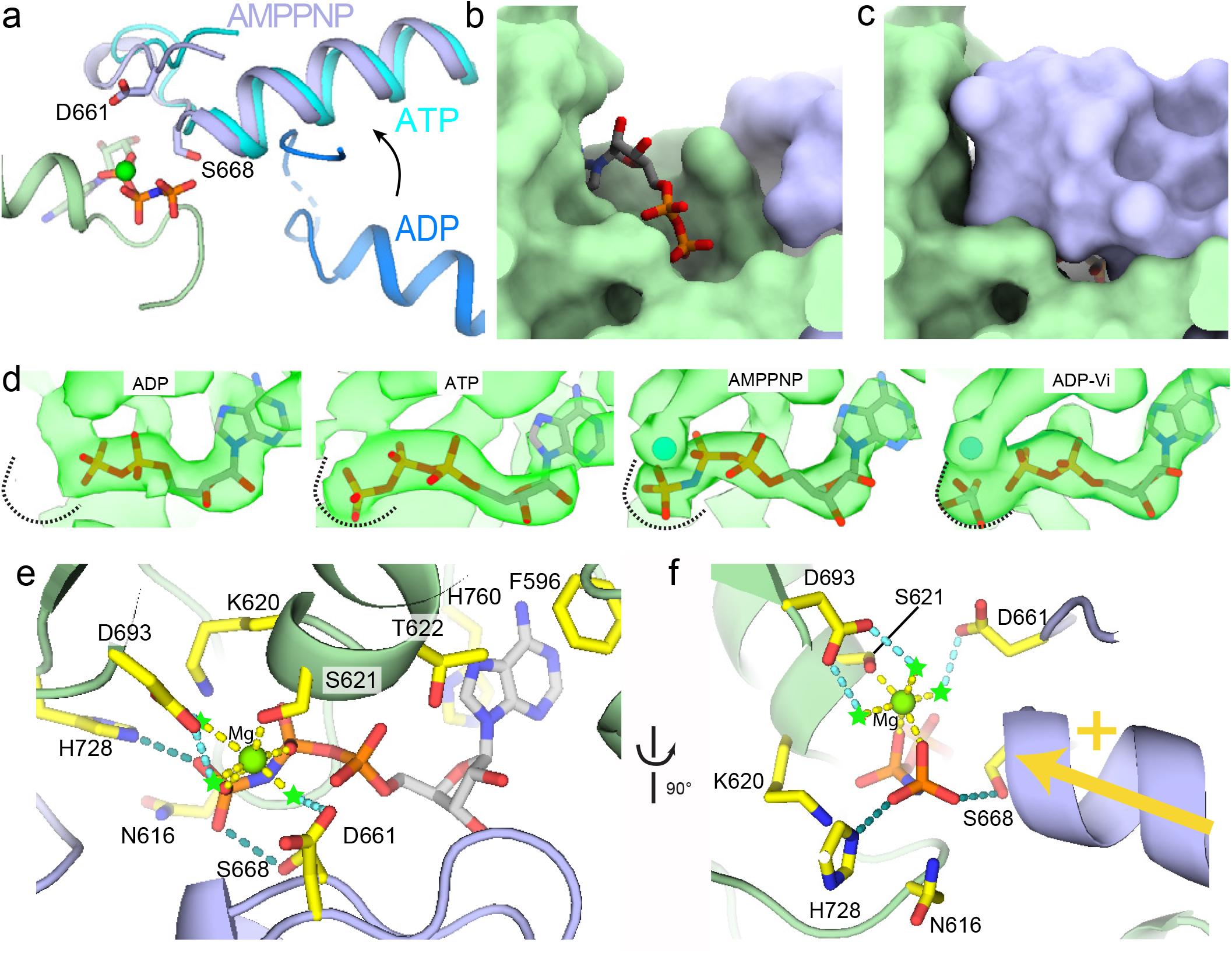
Closing of the MutS dimer completes the ATPase active site. **a**, Movement of the signature loop and helix of monomer B towards the nucleotide in monomer A. Monomer A in green, with AMPPNP and residues Asp661 and Ser668 shown as sticks. Monomer B in dark blue (ADP-bound), light blue (ATP-bound), and pale blue (AMPPNP-bound). **b**, Surface represenation of nucleotide binding site of ADP_2_-bound MutS. Monomer A in green, monomer B in blue, and ADP molecule shown in sticks. **c**, Same representation for ATP:ADP-bound MutS. **d**, Close up of the cryo-EM maps at the nucleotide binding sites in ADP_2_-, ATP:ADP-, AMPPNP_2_-, and ADP-Vi_2_-bound state. The dashed curved line show the boundary of the electron density map in the ADP-Vi_2_-bound state. **e**, Close up of the composite ATP binding site showing the interactions with AMPPNP. Interacting residues are shown in yellow sticks and hydrogen bonds shown as dashed lines. Green stars mark predicted position of water molecules **f**, 90° rotation of the view shown in panel e.

In the ATP-bound site of the ATP:ADP structure the signature motif (GxSTF, residues 666-670) ^29^ of the opposing monomer is further away from the tri-phosphate tail when compared to the structure with AMPPNP and ADP-Vi (Fig. 2a). In addition, only weak density for Mg^2+^ is observed, indicative of a low occupancy (Fig. 2d). In contrast, in the structures of AMPPNP_2_/ADP-Vi_2_-bound MutS the signature loop has moved closer to the phosphate tail of the nucleotide in the opposing monomer and aids in the coordination of the magnesium ion that shows well-defined density (Fig. 2d-f). These two states of the nucleotide binding site (e.g., a sub-optimal conformation in the ATP:ADP state and an optimal conformation in the AMPPNP_2_ and ADP-Vi_2_ state) may be the reason for the bi-phasic ATP hydrolysis rate observed in stop-flow experiments on *E. coli* and *Thermus aquaticus* (Taq) MutS. Here an initial burst of ATPase activity is followed by a slower steady state activity ^30,31^. It is possible that these two rates represent an initial double ATP occupancy of the nucleotide binding sites that are optimal for hydrolysis, followed by a slower rate that may be caused by an asymmetric occupancy (ADP:ATP) that is less optimal for hydrolysis and dictated by the alternating ATPase domains _32_.

The active site conformation of the AMPPNP_2_ and ADP-Vi_2_ structures are identical except for the position of the γ-phosphate that in the ADP-Vi_2_ structure is replaced by the VO_4_^3-^ ion that adopts a pentavalent coordination and has moved away from the β-phosphate, mimicking a post-hydrolysis transition state (Fig. 2d). As the AMPPNP_2_/ADP-Vi_2_ structures represent the hydrolysis competent form, we will discuss the nucleotide binding site in these structure in more detail. The adenosine moiety of the nucleotide is held in place by Phe596 of monomer A (Phe596^A^) and His760^A^ that stack on the base, while Asp616^A^, Lys620^A^ and Thr622^A^ of the Walker A motif (P-loop, residues 614-622) bind the phosphate tail. The Mg^2+^ is coordinated by oxygens of the β- and γ-phosphate of ATP^A^ and Ser621^A^. The three water molecules that complete the octahedral coordination of Mg^2+^ that are observed in the crystal structure of MutS bound to a mismatch and ATP ^33^ are not defined in the cryo-EM maps but are compatible with the position of Asp693^A^ and Asp661 from monomer B (Asp661^B^, Fig. 2e-f). Asp661^B^ can complement the nucleotide binding site of monomer A due to the closing of the MutS dimer that also brings Ser668^B^ in close proximity of the γ-phosphate of ATP^A^. Ser668 is part of the signature motif found in ABC ATPases (sequence GxSTF ^29^) and located immediately upstream of a helix that is pointed with its N-terminus towards the nucleotide in the opposite monomer. This helix deposits the positive charge of its helix dipole onto the negatively charged phosphate tail of the nucleotide through Ser668^B^ (Fig. 2f). With the reduced charge on the phosphate tail and the proper coordination of the Mg^2+^, an activated water molecule can perform a nucleophilic attack on the γ-phosphate. It has been proposed that in the ABC ATPase Rad50, the equivalent of histidine 728^A^ can activate a water molecule to perform the nucleophilic attack on the γ-phosphate ^34^. This was corroborated by recent quantum mechanical and molecular mechanical modelling of the ABC transporter HlyB ^35^. Accordingly, in our structure His728 is well positioned to activate a water molecule as it is pointing towards the γ-phosphate of AMPPNP. Mutation of any of the residues in the ATP active site described above results in a reduction or loss of ATPase activity (see Table 2).

**Table 2.** Effect of mutations on the ATPase activity of *E. coli* MutS.

| **Mutation** | **Reduction (fold)** | **Reference** |
| --- | --- | --- |
| Phe596Ala | 2 | Yang 2001 |
| Lys620Ala | 26 | Haber 1991 |
| Asp661Ala | 1 | Acharya 2008 |
| Ser668Ala | 4 | Lamers 2004 |
| Asn616Ala | 5 | Lamers 2004 |
| His728Ala | 25 | Lamers 2004 |
| Asp693Ala | 22 | Yang 2001 |

Reduction in ATPase activity compared to wild type MutS.

### Two ATP molecules are needed to bring about clamp-state MutS

The structures of AMPPNP_2_-bound and ADP-Vi_2_-bound MutS show a large conformational change of the mismatch and connector domain in one of the monomers (Fig. 1n) that is identical to the conformational change observed in clamp-state MutS ^22,36^. In the ATP:ADP-bound structure however, this conformational change is absent suggesting that a single ATP bound in the MutS dimer is not sufficient to generate enough force to displace the mismatch and connector domains, which is consistent with the observation that in yeast MSH2-MHS6, both monomers need to bind ATP to transform it into a DNA sliding clamp ^37^. To determine if this is indeed the case, we measured the Förster resonance energy transfer (FRET) signal between two fluorophores located on the connector domain (residue 246) of one monomer and on the ATPase domain (residue 798) of the opposing monomer (Fig. 3). Rotation of the connector domain reduces the distance between these two residues and consequently will increase the energy transfer between the two fluorophores. In the presence of ADP, we observe a low FRET signal as predicted from the distance in the ADP-bound structure (Fig 3). Similarly, in the presence of ATP there is no increase in the FRET signal. In contrast, using the non-hydrolysable analogue AMPPNP we observe an increase in the FRET signal as expected from the predicted rotation of the mismatch and connector domain observed in the AMPPNP-bound structures.

**Fig. 3.**
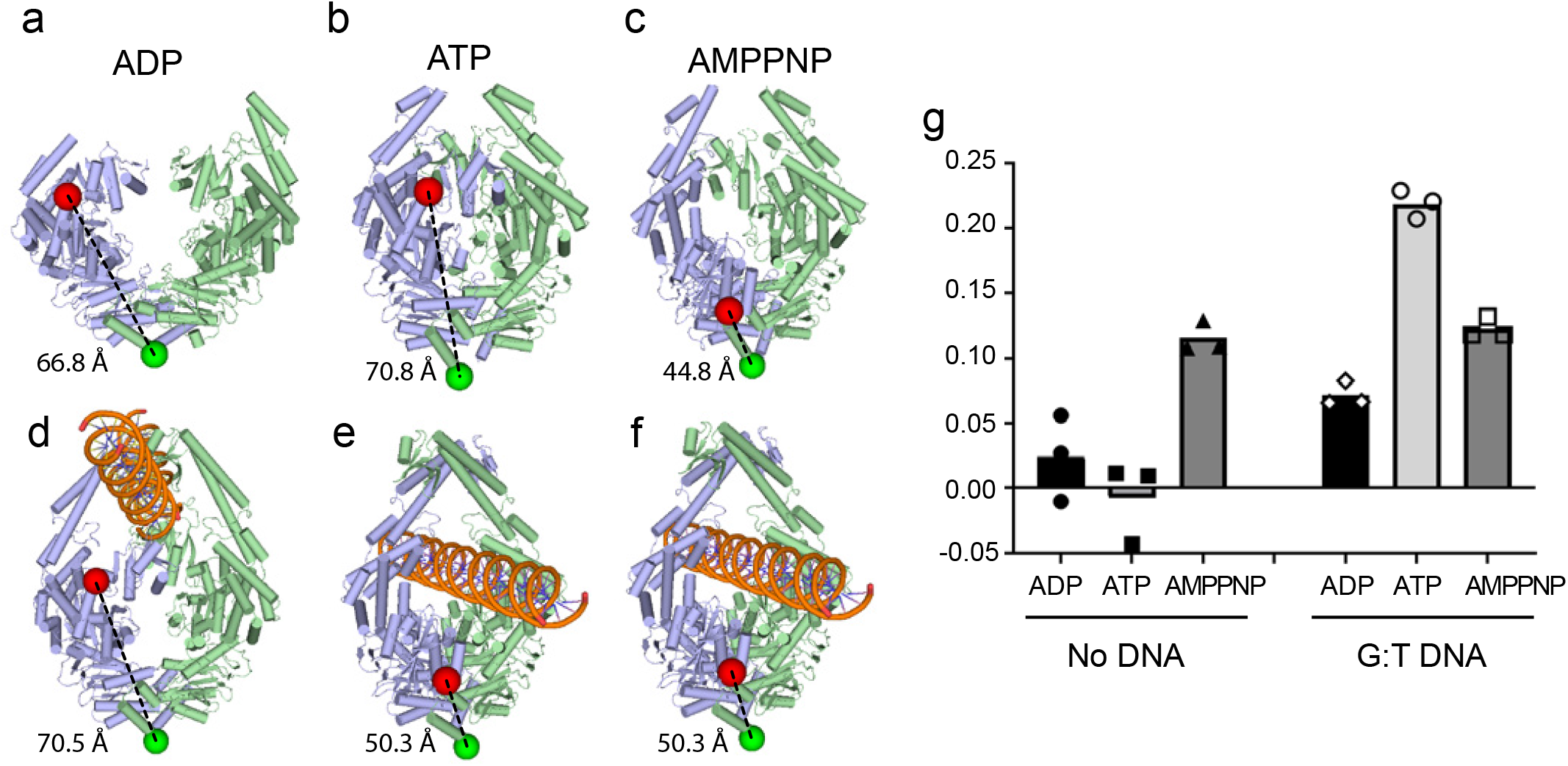
Two ATPs are needed for the MutS clamp formation. **a-c**, Structures of ADP_2_, ATP:ADP, and AMP-PNP_2_-bound MutS. Position of residues 246 and 798 are shown in red and green spheres, respectivily. **d-f**, Structures of DNA-bound MutS in presence of ADP, ATP, and AMPPNP ^22^. **g**, Förster Resonance Energy Transfer (FRET) values between flurorescently labeled residues 246 and 798 in the presence of different nucleotides and DNA. Higher FRET values indicate a closer distance.

In the presence of mismatched DNA and ADP, we observe an increase in the FRET signal, possibly due the reduced flexibility of MutS on DNA, compared to apo MutS that is highly flexible in absence of DNA (Fig. 1i). In the presence of mismatched DNA and ATP, we observe a strong increase of the FRET signal, brought about by the formation of clamp state MutS in which the mismatch and connector domains of both monomers have moved closer to the ATPase domains ^22,25^. In the presence of DNA and AMPPNP we also observe an increase in the FRET signal, although not as strong, possibly due to DNA-free MutS dimers that in the presence of AMPPNP are prevented from binding DNA (Fig. 1n).

Taken together, these results indicate that in the absence of DNA during steady state hydrolysis the MutS dimer binds only one ATP, consistent with the alternating ATPase domains of MutS that have been observed in multiple species ^32,38,39^ and the slow steady state ATP hydrolysis rate ^40^. The single ATP occupancy in the MutS dimer is not enough to bring about a conformational change. This is also observed in mismatch bound MutS that can exist with a single nucleotide triphosphate bound, but not with two ^41^. Concurrently, in the cryo-EM structure of clamp-state MutS, both monomers contain nucleotide triphosphate in their nucleotide binding site ^22^. Moreover, the binding to DNA may actually promote the binding two ATP molecules that are needed to induce the conformational change into the clamp state: the structures of MutS bound to homoduplex and mismatched DNA show the open conformation of the ATPase domains ^22^ that allows for the free exchange with ATP, giving MutS the possibility to load two ATP molecules. This is consistent with the fast release of ADP from mismatch-bound MutS, but not from free MutS ^17,42^

### In clamp state MutS, dsDNA prevents nucleotide release from the ATPase active site

Upon mismatch and ATP binding, MutS transform into the sliding clamp, that remains trapped on the DNA but can freely slide on the DNA until it reaches a stretch of ssDNA or a double strand break where it can dissociate ^18–21^. How MutS is able to remain in the clamp state on DNA for prolonged periods is not known. One possibility is that ATP cannot be hydrolysed in the clamp state, thus keeping MutS in a closed state. However, comparison of the recent structure of clamp state MutS on DNA ^22^ reveals that the position of the ATPase domains is identical to that found in our ADP-Vi_2_ structure, which mimics a post-hydrolysis transition state (Extended Data Fig. 4f). This implies that clamp-state MutS is able to hydrolyse and that therefore it is not ATP that keeps the dimer clamped around the DNA, in agreement with recent observation that Taq MutS can hydrolyse ATP in the clamp state ^43^. Instead, it appears that it is the DNA that keeps the dimer closed. In the sliding clamp structure, the lever domains of the two monomers form an arc that surrounds the DNA duplex and interact with the DNA backbone through numerous residues with a footprint of ∼ 10 base pairs (Fig. 4a-b). In this conformation the ATPase domains are closed, and the nucleotides trapped in the nucleotide binding sites (Fig. 2c). Correspondingly, clamp-state MutS can only release the DNA at open ends or single stranded DNA ^18–21^. Large stretches of ssDNA are produced during downstream events of the mismatch repair cascade through the action of A DNA helicase and DNA exonucleases, which are likely to be encountered by clamp-state MutS, in contrast to double strand breaks, that are not known to play a role in mismatch repair. To measure the minimal length of the ssDNA gap needed to release clamp-state MutS from DNA, we used bio-layer interferometry to measure the release from a DNA substrate containing a mismatch and different sizes of ssDNA gaps in presence of 2 mM ATP (Fig. 4c). The dissociation rate of MutS clamps increases with the size of the single stranded gap, from 0.02 s^-1^ on dsDNA or nicked DNA until it reaches a plateau at 0.08 s^-1^ at a gap of 10 nucleotides or longer (Fig. 4d-e). The ten-nucleotide gap is equal in size to the DNA footprint of the MutS dimer (Fig. 4b), fitting with the notion that as the interactions with the dsDNA are lost, the dimer is no longer able to remain clamped around the DNA and consequently falls off. Hence, by generating ssDNA gaps, UvrD and exonuclease do not only remove the mismatch, but also generate a simple release mechanism for MutS that would otherwise remain trapped on the DNA.

**Fig. 4.**
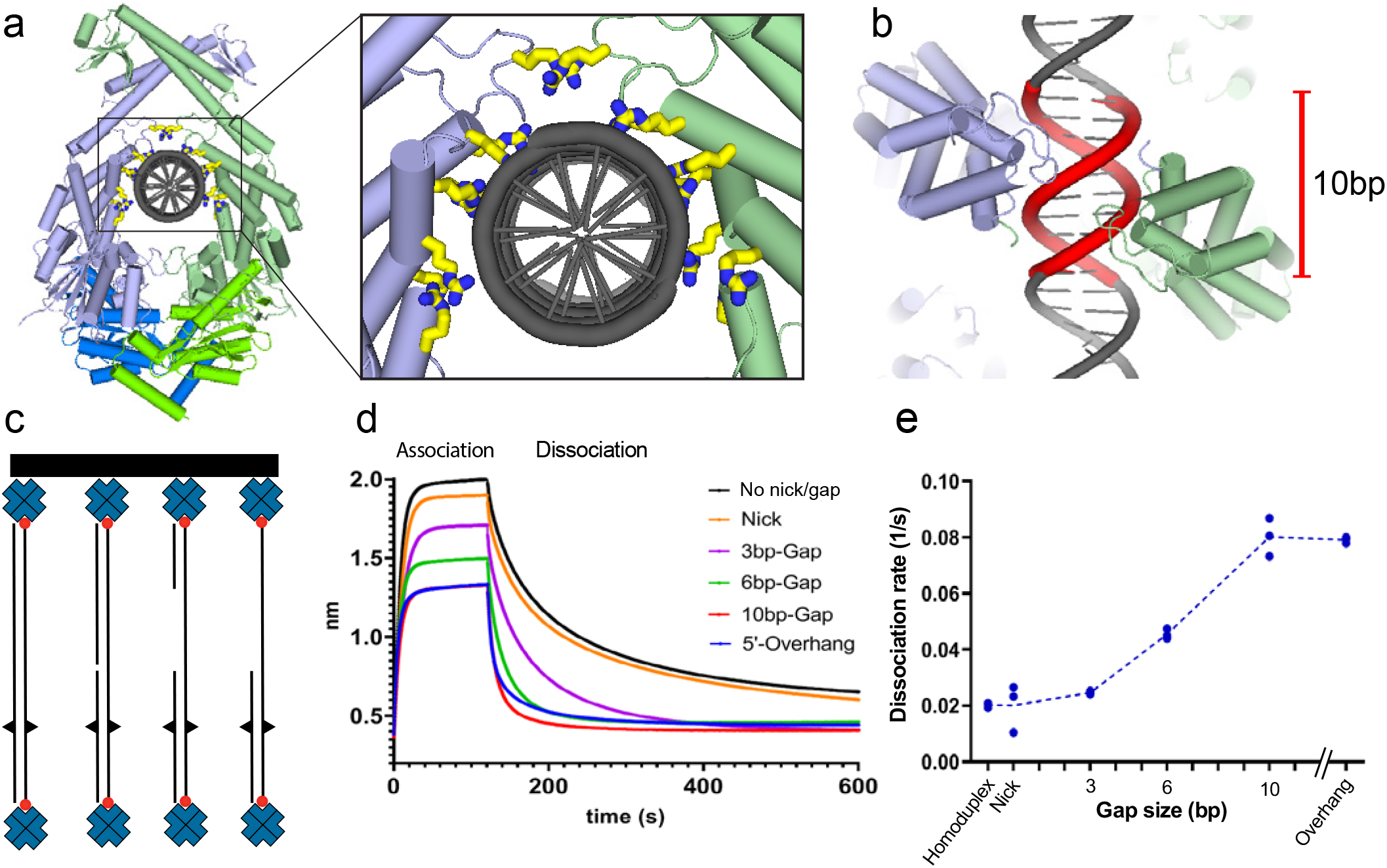
Release of MutS from DNA at a single stranded gap. **a**, Close-up of the protein-DNA interaction in the clamp state of MutS. Arginines and lysines that contact the DNA are shown in yellow sticks. **b**, Footprint of clamp-state MutS dimer on DNA covers 10 base pairs. **c**, Schematic representation of different DNA oligos used in the DNA release experiment. **d**, Association and dissociation of MutS on different DNA substrate in the presence of 2 mM ATP as measured by bio-layer interferometry. **e**, Dissocation rates derived from the binding curves shown in panel d. Dots mark rates calculated from three independent experiments.

## Discussion

### DNA and ATP work together to coordinate the sequential steps of the repair cascade

The four structures presented here provide a detailed view on the mechanism of ATP hydrolysis and how the opening and closing of the dimer is at the core of the catalytic activity. Combined with our recent structures of DNA-bound MutS during different stages of the repair process ^22^, these new structures reveal how ATP hydrolysis and DNA binding co-operate to coordinate the sequential steps of the mismatch repair pathway. In the absence of DNA, the two MutS monomers freely open and close during ATP binding and hydrolysis (Fig. 5a). Due to the alternating ATPase domains of MutS, only one ATP is bound by the dimer, preventing it from assuming the most compact form, which is only reached with the non-hydrolysable ATP analogues AMPPNP or ATPγS. When MutS associates with homoduplex DNA in search for a mismatch, the DNA holds the dimer in an open form thus preventing ATP hydrolysis (Fig. 5b). In this open conformation, the two nucleotide binding sites are freely accessible enabling the loading of two ATP molecules. This open conformation of the ATPase domains is retained when MutS first encounters a mismatch. However, mismatch binding induces a small rearrangement of the clamp domains that acts as a licensing step that enable MutS to transform into the sliding clamp ^22^ but not before both monomers bind ATP.

**Fig. 5.**
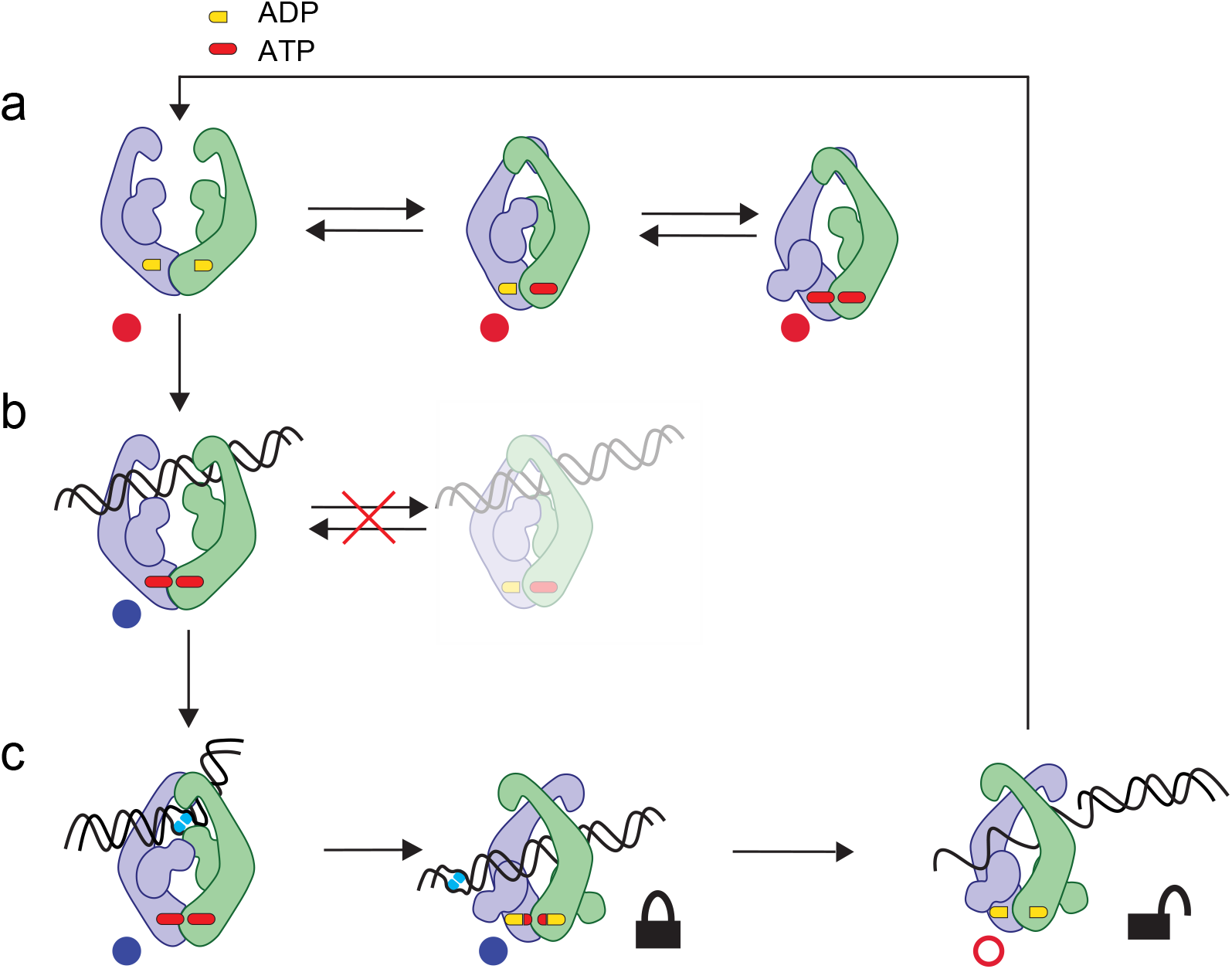
ATP and DNA co-operate to create sequential steps of the repair cascade. Cartoon style respresentation of the different steps of the DNA mismatch repair cascade derived from current and recent cryo-EM work. Red dots indicate structures presented in this work. Blue dots indicate cryo-EM structures presented in previous work ^22^. Red circle indicate predicted structure based on data presented in Fig. 4. **a**, In absence of DNA, the two monomers can freely close and open upon ATP binding and hydrolysis. **b**, When bound to homoduplex DNA the MutS dimer can no longer close and is thereofore not able to hydrolyse ATP (indicated with faded cartoon). The open form of MutS on DNA can freely exchange ADP for ATP. **c**, Upon mismatch binding, MutS transforms into a sliding clamp that can hydrolyse ATP but is locked by double stranded DNA. Only upon encountering single stranded DNA, MutS can dissociate from the DNA and return to the open, ADP-bound form that can re-bind a new DNA substrate.

This transformation into the clamp-state involves a ∼160° rotation of the mismatch and connector domains of both monomers a ∼ 20 Å translation of the DNA towards the centre of the dimer as the lever domains cross each other firmly locking the DNA in the MutS clamp (Fig. 5c) ^22^. In the clamp state, the ATPase active sites are compatible with ATP hydrolysis, yet due to the interactions between the MutS lever domains, the two monomers cannot open up, presumably keeping the ADP-Pi trapped in the nucleotide binding site and preventing the ATPase cycle to continue. Only when MutS encounters ssDNA, generated by UvrD and 5’-3’ or 3’-5’ exonucleases during excision of the newly synthesized strand it is able to release itself from the DNA and resume the hydrolysis cycle.

Thus, as the repair process continues, the size of the ssDNA is increased towards the mismatch, reducing the time that MutS molecules reside on the DNA, until the mismatch is removed and no more loading of MutS molecules takes place. Due to the low processivity of UvrD in the absence of MutS and MutL, the excision of the DNA will soon cease after the removal of a mismatch leaving the DNA open to be resynthesized by a DNA polymerase, completing the repair process.

## Acknowledgement

We thank staff at the LUMC EM facility and The Netherlands Center for Electron Nanoscopy (NeCEN) for help with data collection and data processing. We thank Rafael Fernandez-Leiro for advice on data processing. This work has been supported by a LUMC Research Fellowship to M.H.L., a European Community’s Horizon2020 Marie Skłodowska Curie grant (722433) to P.F., and a European Community’s Horizon2020 Innovative Training Network Grant to A.B. and V.K.. Access to NeCEN was supported by the Netherlands Electron Microscopy Infrastructure (NEMI), project 184.034.014 of the National Roadmap for Large-Scale Research Infrastructure of the Dutch Research Council (NWO).

## Author contributions

M.H.L. and A.B. conceived the overall experimental design; A.B. prepared samples, collected and processed cryo-EM data; A.B. purified proteins and performed BLI experiments. V.K. performed FRET experiments; A.B. and M.H.L. wrote the manuscript with contributions from all authors.

## Methods

### Materials

All chemicals were purchased from Sigma Aldrich, unless indicated otherwise. All chromatography columns were purchased from Cytiva.

### Protein Expression and Purification

The dimeric version of *E. coli* MutS, R840E ^36^ was cloned into vector pETNKI-his3C-LIC-amp ^44^. Plasmids were transformed in *E. coli* BL21(DE3) pLysS cells and plated onto LB-agar with 50 μg/ml chloramphenicol and 100 μg/ml ampicillin. All the colonies were scraped from the plate and distributed over 6 x 500 ml Terrific Broth supplemented with 50 μg/ml chloramphenicol, 100 μg/ml ampicillin, 1 mM MgCl_2_ and 1% glucose. Cells were grown at 37°C to OD_600_ ∼7, diluted with 1 volume Terrific Broth and induced with a final concentration of 1 mM isopropyl 1-thio-β-D-galactopyranoside (IPTG) for 2 hrs at 30 °C. The proteins were purified using a modified procedure from ^45^. All subsequent steps were performed using basic buffer (25 mM Hepes pH 7.5, 5mM MgCl2, 2 mM DTT) supplemented with NaCl or Imidazole as indicated.

Harvested cells were resuspended in basic buffer with 500 mM NaCl and 10 mM imidazole and lysed by sonication, followed by centrifugation at 14k x g. The supernatant was injected onto a 5 ml HisTrap HP column and eluted with a gradient to 500 mM Imidazole in basic buffer with 500 mM NaCl. Pooled fractions where diluted in ten volumes of basic buffer, injected onto 5 ml HiTrap Heparin column and eluted with a gradient to 1 M NaCl in basic buffer. Pooled fractions were concentrated and injected into a gel filtration column Superdex 200 16/600 equilibrated in basic buffer with 100 mM NaCl. Pooled fractions were injected onto a 5 ml HiTrap Q column equilibrated in the same buffer and eluted with a gradient to 1M NaCl in basic buffer.

### Cryo-EM sample preparation and imaging

Purified MutS was diluted to 4 µM in 20 mM Tris pH 8.5, NaCl 150 mM, 5mM MgCl2, 2 mM DTT and 0.01% (w/v) Tween 20. The diluted protein was incubated with 3 mM of either ADP, ATP, AMPPNP or ADP-Vi. Cu R1.2/1.3 or R2/1 holey carbon grids (Quantifoil, Groslöbichau, Germany) were glow discharged at 25 mA for 45 seconds using an Emitech K950 apparatus (Quorum, Laughton, United Kingdom). 3 μl of MutS-nucleotide sample were adsorbed onto glow-discharged grids and blotted for 1 second at 80% humidity at 4°C and flash frozen in liquid ethane using an EM GP plunge freezer (Leica). All cryo-EM data was collected at the The Netherlands Center for Electron Nanoscopy (NeCEN). The grids were loaded into a Titan Krios (FEI) electron microscope operating at 300 kV with a K2 or K3 direct electron detector equipped with a Bioquantum energy filter (Gatan, Pleasonton USA). The slit width of the energy filter was set to 20 eV. Images were recorded with EPU software (Thermo Fisher Scientific) in counting mode. Dose, magnification and effective pixel size are detailed in Table 1.

### Cryo-EM image processing

All image processing was performed using Relion 3.1 ^46^. The images were drift corrected using Relion’s own (CPU-based) implementation of the UCSF Motioncor2 program, and defocus was estimated using gCTF ^47^. LoG-based auto-picking was performed on a subset of micrographs and picked particles where 2D classified. Selected classes from the 2D classification were used as references to autopick particles from the full data sets. After two or three rounds of 2D classification, classes with different orientations were selected for initial model generation in Relion. The initial model was used as reference for 3D classification into different classes. The selected classes from 3D classification were subjected to 3D auto refinement. The defocus values were further refined using CTF Refinement in Relion followed by Bayesian polishing. Another round of 3D auto refinement was performed on these polished particles. The density for MutS ADP-Vi was improved by using focused classification without image alignment. All maps were post-processed to correct for modulation transfer function of the detector and sharpened by applying a negative B factor, as determined automatically by Relion. A soft mask was applied during post processing to generate FSC curves to yield maps of average resolutions of 3.4 Å for MutS AMPPNP, 4.8 Å for MutS ADP, 3.8 Å for MutS ADP-Vi and 3.3 Å for MutS ATP.

Model building was performed using Coot ^48^, REFMAC5 ^49^, the CCPEM-suite ^50^ and Phenix ^51^. For the ATP:ADP, AMPPNP_2_ and ADPVi_2_ structures we could build detailed model into the high-resolution maps, while for the ADP_2_ was limited to rigid body fitting of existing structures and restrained refinement with REFMAC. Details on model refinement and validation are in Table 1. In brief, model building started by rigid-body fitting known crystal structures (PDB 1E3M and PDB 5AKB) into the different maps using Coot. Next, we carried out one round of refinement in Refmac5 using jelly-body restraints, and the model was further adjusted in Coot. After initial refinement, we generated improved-resolution EM maps using the SuperEM method ^52^, which aided in model building and refinement. A final refinement round and validation of the model and data were carried out using Refmac5 with proSmart restraints within the CCPEM suite, with additional model validation using MolProbity within Phenix.

### Negative stain EM sample preparation and imaging

Purified MutS was diluted to 0.04 μM in 20 mM Tris pH 8.5, NaCl 150 mM, 5mM MgCl2, 2mM DTT, 0.01% Tween20 and 1 mM ADP. Grids were glow discharged 60 seconds at 25 mA using an EMITECH K950 apparatus. Three μl of sample were absorbed to glow-discharged carbon-coated copper grids (C-Flat200-CU) and incubated for 30 s, before being partially blotted. The grids were sequentially stained with three μl of 2.3% uranyl acetate, incubated for 30 s and then blotted dry. Micrographs were collected on an FEI Tecnai Biotwin electron microscope equipped with a LaB6 filament and operated at 120 kV and 49,000× nominal magnification. The total dose was 20 e−/Å2 and the defocus values ranged from −0.5 to −3.0 μm. Images were collected with a FEI Eagle 4k x 4K CCD camera with final pixel size of 4.44 Å.

### Negative stain image processing

All image processing was performed using Relion 3.1 ^46^. Contrast transfer function estimation was performed with CTFFIND4.1 ^53^. A small set of particles were manually picked, 2D classified and used as references for the auto picking procedure. 7845 particles were extracted from a total of 73 micrographs. The extraction procedure was performed without inverting contrast. Extracted particles were subjected to reference-free 2D average without performing CTF correction.

### Biolayer Interferometry DNA substrates

Six different DNA substrates with different gap lengths were prepared for the assays (Supplementary Table 1). The substrates were prepared as follows: monovalent streptavidin and a 5’ biotinylated primer were mixed in a 1:1 ratio at a nominal concentration of 20 μM and purified via analytical gel filtration using a Superdex 200 increase (3.2/30) column (Cytiva) equilibrated in Tris 10 mM pH 8.0, NaCl 50 mM, EDTA 1mM. Fractions containing biotinylated primer were mixed with the 5’ biotinylated DNA template to create the final DNA substrate. The efficient annealing of primer and template was confirmed with a 15% polyacrylamide 0.5 x Tris Borate EDTA gel. Annealed substrates were stored at −20°C at a concentration of 5 μM.

### Biolayer Interferometry DNA assay

Association and dissociation kinetics between MutS and DNA were observed using the Octet RED96 (Sartorius, Göttingen, Germany). All interactions studies were performed with Streptavidin biosensors (Sartorius, Göttingen, Germany) conjugated to the biotinylate DNA substrates. All the experiments were performed in the following buffer: Hepes 25 mM pH 7.5, NaCl 150 mM, MgCl2 5mM, DTT 2mM, BSA 0.5 mg/ml, Tween 20 0.01%. The DNA substrates were added in the loading step at 100 nM, until the threshold value of 0.36 nm was reached. The association step was performed with MutS 200 nM in presence of 2 mM ATP. The dissociation step was performed in buffer with 2mM ATP. Kinetic analysis were performed using the Octet Data Analysis software package version 7.1.

### FRET assay

Förster Resonance Energy Transfer (FRET) between labelled connector domain of one subunit and the ATPase domain of the other subunit was measured using heterodimeric MutS dimers similar as described before ^36^. For labelling, proteins were diluted to 40 μM in 150 µl of buffer (10 mM HEPES/KOH (pH 8.0), 200 mM KCl and 1 mM EDTA) and labelled with a 5-fold molar excess of Alexa Fluor 488 maleimide or Alexa Fluor 647 maleimide (Invitrogen, Thermo Fisher Scientific, Waltham, MA) for 2 hours on ice in the dark according to the manufacturer’s instruction. Excessive dye was removed using Zeba Spin Desalting columns (Thermo Fisher Scientific, Waltham, MA). The degree of labelling was determined from the absorbance spectra recorded from 220–700 nm (nanodrop) according to the manufactures instructions as described previously ^36^. Heterodimers were allowed to form by mixing 10 µM of MutS798C labelled with Alexa Fluor 488 and MutS246C labelled with Alexa Fluor 647, and incubation on ice for at least 1.5 hours in the absence of nucleotides in buffer (25 mM HEPES-KOH pH 7.5, 5 mM MgCl_2_, 150 mM KCl, and 0.05 % (v/v) Tween 20). Aliquots of the reaction were flash-frozen in liquid nitrogen and stored at −80 °C. A 30-bp DNA substrate with a central G:T mismatch was prepared by annealing T-strand (5’ dig-AATTGCACCGAGCT<u>T</u>GATCCTCGATGATCC-dig 3’) with G-strand (5’ GGATCATCGAG<u>G</u>ATCGAGCTCGGTGCAATT 3’). Underlines mark the G:T mismatch. The T-strand contains digoxigenin on either side, so that both DNA ends are blocked with 200 nM anti-digoxigenin Fab fragments (Roche Diagnostics, F. Hoffmann-La Roche Ltd, Switzerland) MutS heterodimers (50 nM monomer) were pre-incubated in 200 µl of FB150 buffer (25 mM HEPES-KOH pH 7.5, 5 mM MgCl_2_, 150 mM KCl, and 0.05 % (v/v) Tween 20) in 96-well plates at room temperature for 120 seconds. Next, 50 nM of the G:T DNA substrate was added, followed by addition of 1 mM ATP, ADP, or AMPPNP (Jena Bioscience, Jena, Germany, and AMPPNP from Sigma-Aldrich).

FRET was determined by measuring the fluorescence signals in microplate reader (TECAN infinite F200, Tecan Group Ltd, Switzerland) before and 10 minutes later after the addition of the dNTPs. Fluorescence signals were measured in three channels (donor, acceptor, FRET) with the following filter combinations: (donor ex. 450 nm (width 20 nm) em. 535 nm (width 25 nm), acceptor ex. 620 nm (width 10 nm) em. 670 nm (width 25 nm), FRET ex. 485 nm (width 20 mm) em. 670 nm (width 25 nm) filter.

---

**Extended Data Fig. 1.**
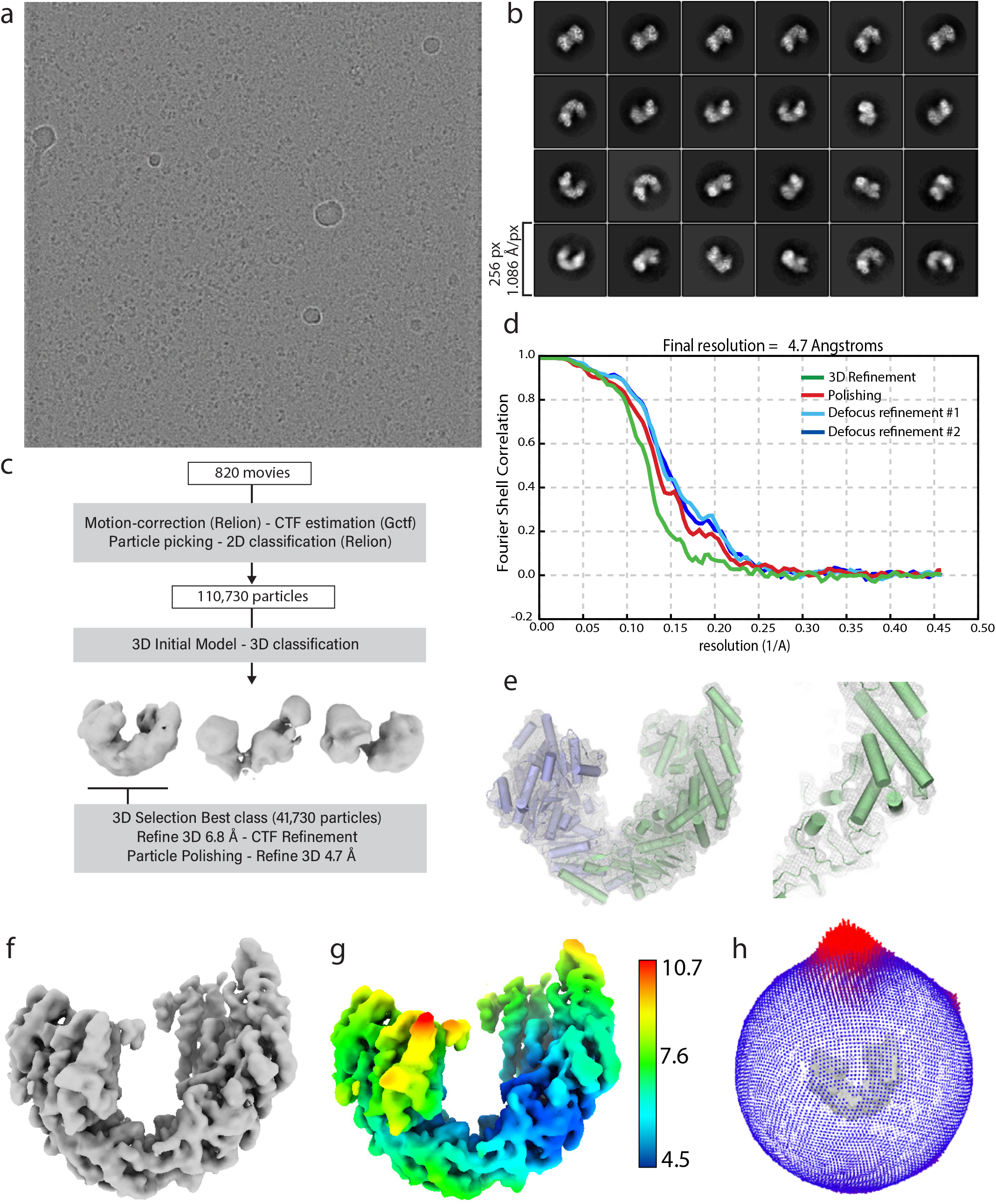
CryoEM data analysis of ADP bound MutS. **a**, Representativ micrograph. **b**, 2D class averages from full dataset. **c**, Schematic representation of main data processing procedures. See methods section for more details. **d**, Fourier Shell Correlation between half-maps from subsequent refinements in the processing procedures. **e**, Detail of model fit to map. **f**, Final map obtained applying SuperEM code to Relion post-processed map. **g**, final map colored by local resolution. **h**, Orientation distribution in final set of refined particles.

**Extended Data Fig. 2.**
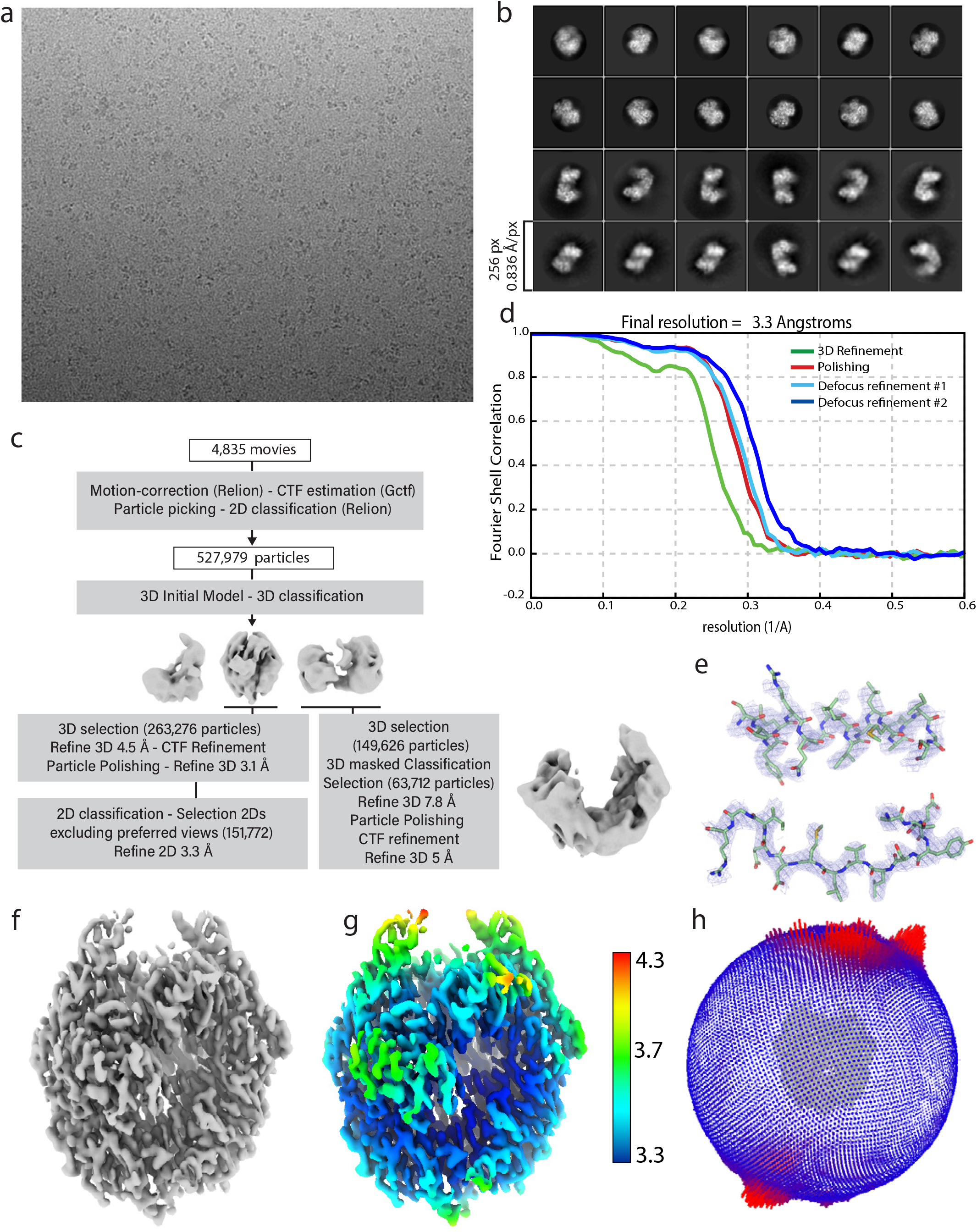
CryoEM data analysis of ADP-ATP bound MutS. **a**, Representativ micrograph. **b**, 2D class averages from full dataset. **c**, Schematic representation of main data processing procedures. See methods section for more details. **d**, Fourier Shell Correlation between half-maps from subsequent refinements in the processing procedures. **e**, Detail of model fit to map. **f**, Final map obtained applying SuperEM code to Relion post-processed map. **g**, final map colored by local resolution. **h**, Orientation distribution in final set of refined particles.

**Extended Data Fig. 3.**
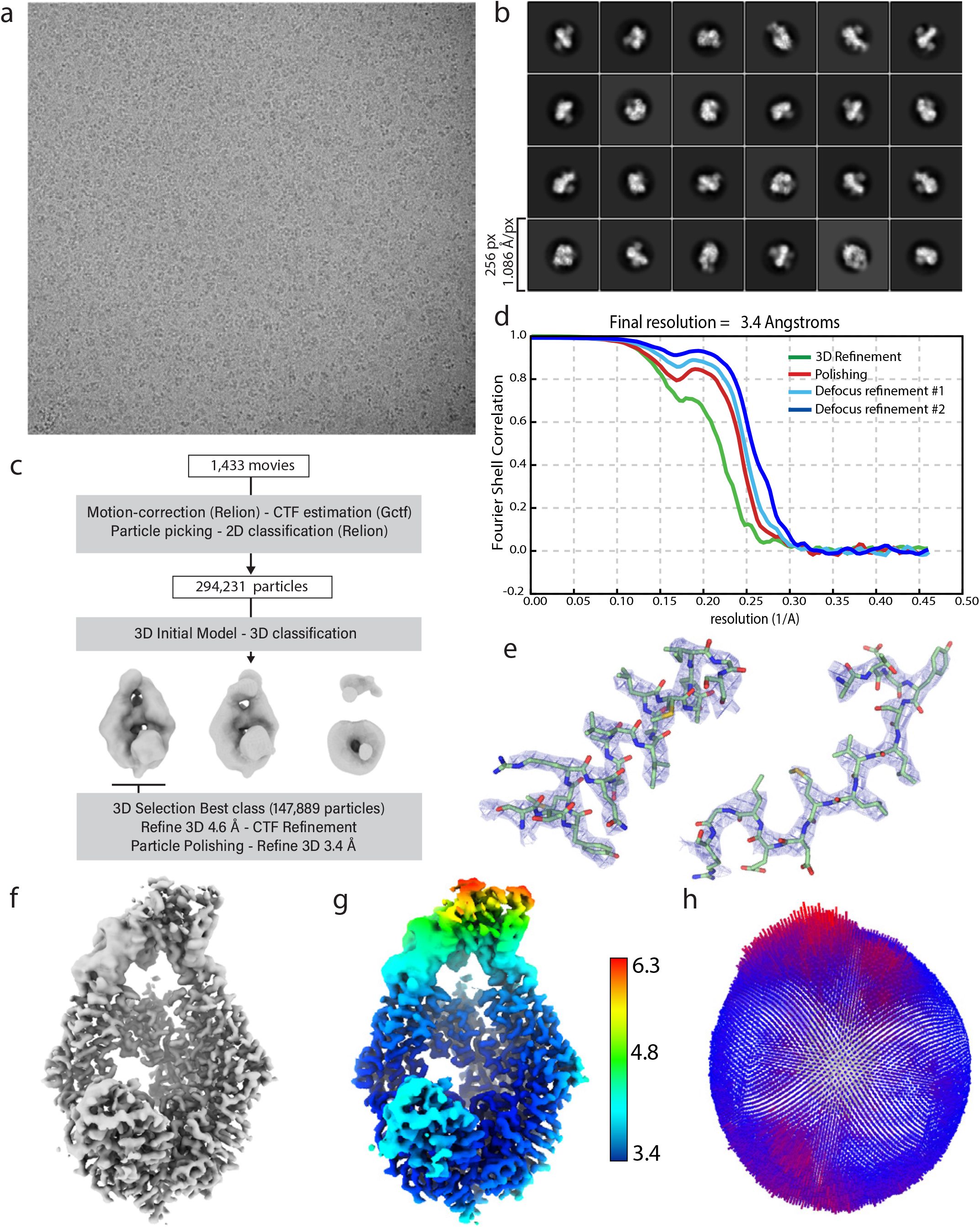
CryoEM data analysis of ANPPNP bound MutS. **a**, Representativ micrograph. **b**, 2D class averages from full dataset. **c**, Schematic representation of main data processing procedures. See methods section for more details. **d**, Fourier Shell Correlation between half-maps from subsequent refinements in the processing procedures. **e**, Detail of model fit to map. **f**, Final map obtained applying SuperEM code to Relion post-processed map. **g**, final map colored by local resolution. **h**, Orientation distribution in final set of refined particles.

**Extended Data Fig. 4.**
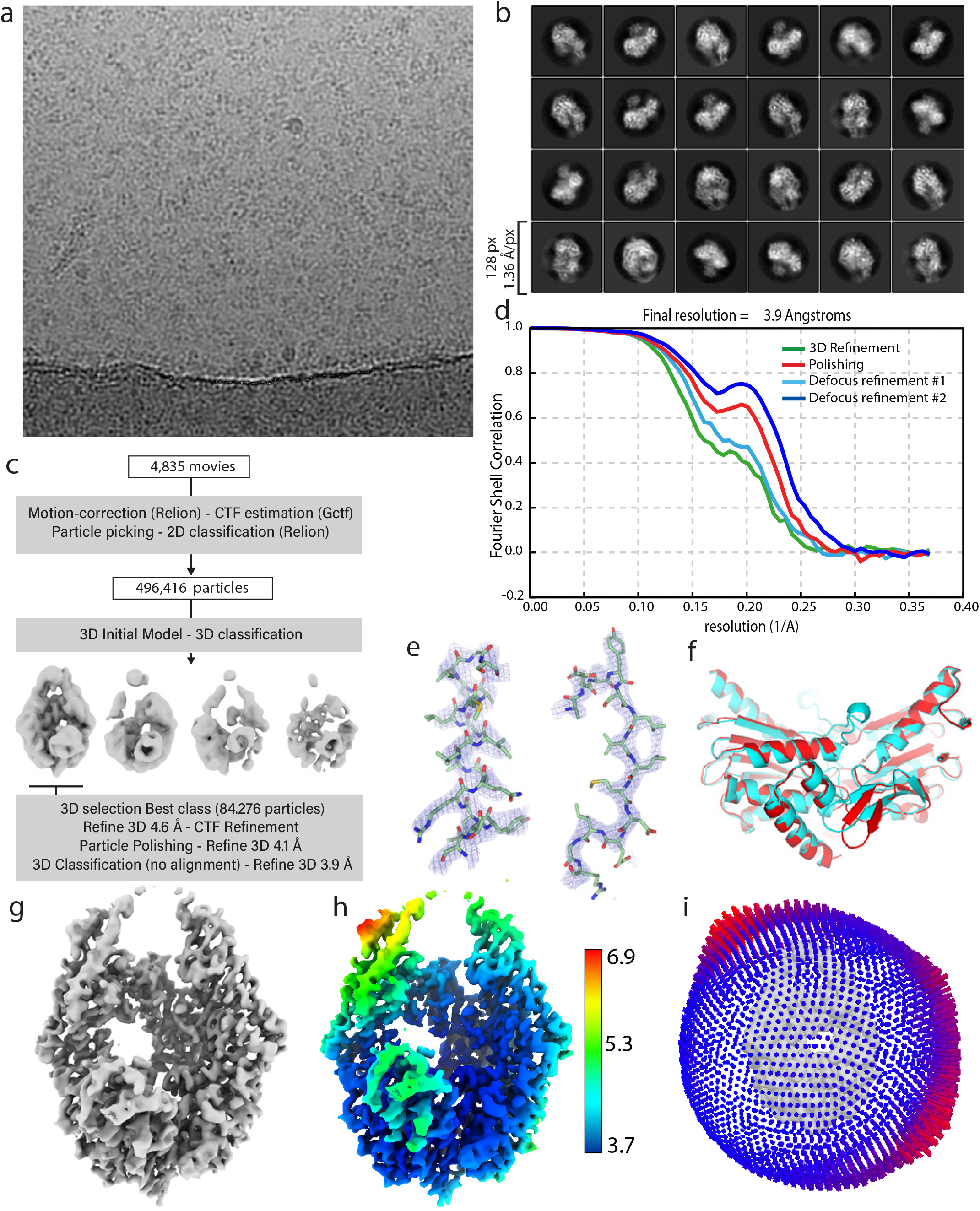
CryoEM data analysis of ADP-Vi bound MutS. **a**, Representativ micrograph. **b**, 2D class averages from full dataset. **c**, Schematic representation of main data processing procedures. See methods section for more details. **d**, Fourier Shell Correlation between half-maps from subsequent refinements in the processing procedures. **e**, Detail of model fit to map. **f**, Superimposition of nucleotide binding domains of MutS in ADP-Vi conformation and MutS in sliding clamp MutL bound conformation (**Fernandez-Leiro 2021**). **g** Final map obtained applying SuperEM code to Relion post-processed map. **h**, final map colored by local resolution. **i**, Orientation distribution in final set of refined particles.

## References

1. Li, Z., Pearlman, A. H. & Hsieh, P. DNA mismatch repair and the DNA damage response. DNA Repair (Amst) 38, 94–101 (2016).

2. Jiricny, J. Postreplicative mismatch repair. Cold Spring Harb Perspect Biol 5, a012633 (2013).

3. Welsh, K. M., Lu, A. L., Clark, S. & Modrich, P. Isolation and characterization of the Escherichia coli mutH gene product. J. Biol. Chem. 262, 15624–15629 (1987).

4. Junop, M. S., Yang, W., Funchain, P., Clendenin, W. & Miller, J. H. In vitro and in vivo studies of MutS, MutL and MutH mutants: correlation of mismatch repair and DNA recombination. DNA Repair (Amst) 2, 387–405 (2003).

5. Hall, M. C. & Matson, S. W. The Escherichia coli MutL protein physically interacts with MutH and stimulates the MutH-associated endonuclease activity. J. Biol. Chem. 274, 1306–1312 (1999).

6. Kadyrov, F. A., Dzantiev, L., Constantin, N. & Modrich, P. Endonucleolytic function of MutLalpha in human mismatch repair. Cell 126, 297–308 (2006).

7. Fukui, K., Nishida, M., Nakagawa, N., Masui, R. & Kuramitsu, S. Bound nucleotide controls the endonuclease activity of mismatch repair enzyme MutL. J. Biol. Chem. 283, 12136–12145 (2008).

8. Pluciennik, A. et al. PCNA function in the activation and strand direction of MutLα endonuclease in mismatch repair. Proc. Natl. Acad. Sci. USA 107, 16066–16071 (2010).

9. Pillon, M. C., Miller, J. H. & Guarné, A. The endonuclease domain of MutL interacts with the β sliding clamp. DNA Repair (Amst) 10, 87–93 (2011).

10. Yang, H., Yung, M., Sikavi, C. & Miller, J. H. The role of Bacillus anthracis RecD2 helicase in DNA mismatch repair. DNA Repair (Amst) 10, 1121–1130 (2011).

11. Walsh, B. W. et al. RecD2 helicase limits replication fork stress in Bacillus subtilis. J. Bacteriol. 196, 1359–1368 (2014).

12. Burdett, V., Baitinger, C., Viswanathan, M., Lovett, S. T. & Modrich, P. In vivo requirement for RecJ, ExoVII, ExoI, and ExoX in methyl-directed mismatch repair. Proc. Natl. Acad. Sci. USA 98, 6765–6770 (2001).

13. Yamaguchi, M., Dao, V. & Modrich, P. MutS and MutL activate DNA helicase II in a mismatch-dependent manner. J. Biol. Chem. 273, 9197–9201 (1998).

14. Lahue, R. S., Au, K. G. & Modrich, P. DNA mismatch correction in a defined system. Science 245, 160–164 (1989).

15. Goellner, E. M., Putnam, C. D. & Kolodner, R. D. Exonuclease 1-dependent and independent mismatch repair. DNA Repair (Amst) 32, 24–32 (2015).

16. Schofield, M. J., Nayak, S., Scott, T. H., Du, C. & Hsieh, P. Interaction of Escherichia coli MutS and MutL at a DNA mismatch. 276, 28291–28299 (2001).

17. Acharya, S., Foster, P. L., Brooks, P. & Fishel, R. The coordinated functions of the E. coli MutS and MutL proteins in mismatch repair. 12, 233–246 (2003).

18. Gradia, S. et al. hMSH2-hMSH6 forms a hydrolysis-independent sliding clamp on mismatched DNA. 3, 255–261 (1999).

19. Blackwell, L. J., Bjornson, K. P., Allen, D. J. & Modrich, P. Distinct MutS DNA-binding modes that are differentially modulated by ATP binding and hydrolysis. 276, 34339–34347 (2001).

20. Heo, S.-D., Cho, M., Ku, J. K. & Ban, C. Steady-state ATPase activity of E. coli MutS modulated by its dissociation from heteroduplex DNA. Biochem Biophys Res Commun 364, 264–269 (2007).

21. Jeong, C. et al. MutS switches between two fundamentally distinct clamps during mismatch repair. Nat. Struct. Mol. Biol. (2011). doi:10.1038/nsmb.2009

22. Fernandez-Leiro, R. et al. The selection process of licensing a DNA mismatch for repair. Nat. Struct. Mol. Biol. 28, 373–381 (2021).

23. Hingorani, M. M. Mismatch binding, ADP–ATP exchange and intramolecular signaling during mismatch repair. DNA Repair (Amst) 38, 24–31 (2016).

24. Davies, D. R. & Hol, W. G. J. The power of vanadate in crystallographic investigations of phosphoryl transfer enzymes. FEBS Lett. 577, 315–321 (2004).

25. Groothuizen, F. S. et al. MutS/MutL crystal structure reveals that the MutS sliding clamp loads MutL onto DNA. Elife 4, e06744 (2015).

26. Hopfner, K.-P. Invited review: Architectures and mechanisms of ATP binding cassette proteins. Biopolymers 105, 492–504 (2016).

27. Hol, W. G. The role of the alpha-helix dipole in protein function and structure. Prog Biophys Mol Biol 45, 149–195 (1985).

28. Hopfner, K.-P. & Tainer, J. A. Rad50/SMC proteins and ABC transporters: unifying concepts from high-resolution structures. Curr Opin Struct Biol 13, 249–255 (2003).

29. Acharya, S. Mutations in the signature motif in MutS affect ATP-induced clamp formation and mismatch repair. Mol Microbiol 69, 1544–1559 (2008).

30. Bjornson, K. P., Allen, D. J. & Modrich, P. Modulation of MutS ATP hydrolysis by DNA cofactors. Biochemistry 39, 3176–3183 (2000).

31. Antony, E. & Hingorani, M. M. Asymmetric ATP binding and hydrolysis activity of the Thermus aquaticus MutS dimer is key to modulation of its interactions with mismatched DNA. 43, 13115–13128 (2004).

32. Lamers, M. H., Winterwerp, H. H. K. & Sixma, T. K. The alternating ATPase domains of MutS control DNA mismatch repair. 22, 746–756 (2003).

33. Lamers, M. H. et al. ATP increases the affinity between MutS ATPase domains. Implications for ATP hydrolysis and conformational changes. 279, 43879–43885 (2004).

34. Hopfner, K. P. et al. Structural biology of Rad50 ATPase: ATP-driven conformational control in DNA double-strand break repair and the ABC-ATPase superfamily. Cell 101, 789–800 (2000).

35. Zhou, Y., Ojeda-May, P. & Pu, J. H-loop histidine catalyzes ATP hydrolysis in the E. coli ABC-transporter HlyB. Phys. Chem. Chem. Phys. 15, 15811–15815 (2013).

36. Groothuizen, F. S. et al. Using stable MutS dimers and tetramers to quantitatively analyze DNA mismatch recognition and sliding clamp formation. Nucleic Acids Res. 41, 8166–8181 (2013).

37. Mazur, D. J., Mendillo, M. L. & Kolodner, R. D. Inhibition of Msh6 ATPase activity by mispaired DNA induces a Msh2(ATP)-Msh6(ATP) state capable of hydrolysis-independent movement along DNA. Mol. Cell 22, 39–49 (2006).

38. Iaccarino, I., Marra, G., Palombo, F. & Jiricny, J. hMSH2 and hMSH6 play distinct roles in mismatch binding and contribute differently to the ATPase activity of hMutSalpha. EMBO J. 17, 2677–2686 (1998).

39. Studamire, B., Quach, T. & Alani, E. Saccharomyces cerevisiae Msh2p and Msh6p ATPase activities are both required during mismatch repair. Mol Cell Biol 18, 7590– 7601 (1998).

40. Sharma, A., Doucette, C., Biro, F. N. & Hingorani, M. M. Slow conformational changes in MutS and DNA direct ordered transitions between mismatch search, recognition and signaling of DNA repair. J. Mol. Biol. 425, 4192–4205 (2013).

41. Monti, M. C. et al. Native mass spectrometry provides direct evidence for DNA mismatch-induced regulation of asymmetric nucleotide binding in mismatch repair protein MutS. Nucleic Acids Res. 39, 8052–8064 (2011).

42. Gradia, S., Acharya, S. & Fishel, R. The human mismatch recognition complex hMSH2-hMSH6 functions as a novel molecular switch. 91, 995–1005 (1997).

43. Hao, P. et al. Recurrent mismatch binding by MutS mobile clamps on DNA localizes repair complexes nearby. Proc. Natl. Acad. Sci. USA 117, 17775–17784 (2020).

44. Luna-Vargas, M. P. A. et al. Enabling high-throughput ligation-independent cloning and protein expression for the family of ubiquitin specific proteases. J. Struct. Biol. 175, 113–119 (2011).

45. Feng, G. & Winkler, M. E. Single-step purifications of His6-MutH, His6-MutL and His6-MutS repair proteins of escherichia coli K-12. BioTechniques 19, 956–965 (1995).

46. Zivanov, J. et al. New tools for automated high-resolution cryo-EM structure determination in RELION-3. Elife 7, 163 (2018).

47. Zhang, K. Gctf: Real-time CTF determination and correction. J. Struct. Biol. 193, 1–12 (2016).

48. Emsley, P., Lohkamp, B., Scott, W. G. & Cowtan, K. Features and development of Coot. Acta Crystallogr. D Biol. Crystallogr. 66, 486–501 (2010).

49. Murshudov, G. N. et al. REFMAC5 for the refinement of macromolecular crystal structures. Acta Crystallogr. D Biol. Crystallogr. 67, 355–367 (2011).

50. Nicholls, R. A., Tykac, M., Kovalevskiy, O. & Murshudov, G. N. Current approaches for the fitting and refinement of atomic models into cryo-EM maps using CCP-EM. Acta Crystallogr D Struct Biol 74, 492–505 (2018).

51. Liebschner, D. et al. Macromolecular structure determination using X-rays, neutrons and electrons: recent developments in Phenix. Acta Crystallogr D Struct Biol 75, 861– 877 (2019).

52. Subramaniya, S. R. M. V., Terashi, G. & Kihara, D. Super-Resolution Cryo-EM Maps With 3D Deep Generative Networks. bioRxiv 1–30 (2021). doi:10.1101/2021.01.12.426430

53. Rohou, A. & Grigorieff, N. CTFFIND4: Fast and accurate defocus estimation from electron micrographs. J. Struct. Biol. 192, 216–221 (2015).

